# mzrtsim: Raw Data Simulation for Reproducible Gas/Liquid Chromatography–Mass Spectrometry Based Non-targeted Metabolomics Data Analysis

**DOI:** 10.1101/2023.11.14.567024

**Authors:** Miao Yu, Vivek Philip

## Abstract

Reproducibility of data analysis is pivotal in the context of non-targeted metabolomics based on mass spectrometry coupled with chromatography. While various algorithms have been proposed for feature or peak extraction, their validation often revolves around a limited set of known compounds or standards. While data simulation is widely used in other omics studies, simulation are focused on the feature level, neglecting uncertainties inherent in the feature or peak extraction process for metabolomics mass spectrometry data. In this technique note, we introduce a R package called ‘mzrtsim’ to simulate gas/liquid chromatography full scan raw data in the mzML format. Unlike simulations solely based on virtual features, our approach leverages experimental spectral data from the MassBank of North America (MoNA) and the human metabolome database (HMDB). We developed algorithms to simulate chromatographic peaks, accounting for the tailing factor. The results of our study demonstrate the potential of this tool for comparing established metabolomics software (e.g., XCMS, mzMine, and OpenMS) against a ground truth. We found that the investigated software introduced false positive peaks and/or loss compounds with less peaks. They also showed different sensitivity to tailing and leading peaks and we also found setting appropriate intensity cutoffs will not lose information for the following statistical analysis. This R package is free and available online (https://github.com/yufree/mzrtsim).

**TOC:** 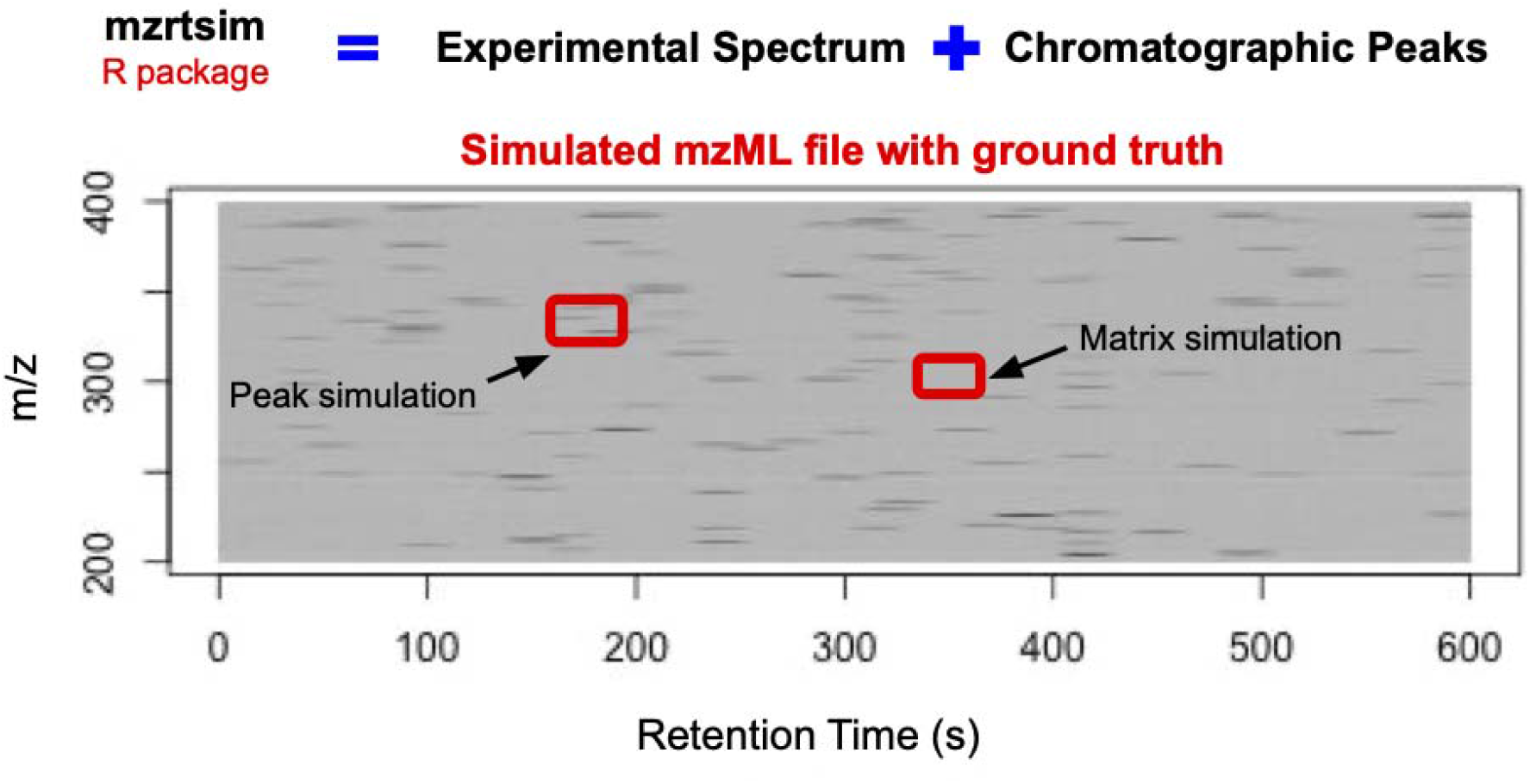

## Introduction

Metabolomics is focused on the comprehensive identification and precise quantification of metabolites within a biological system^1^. To conduct metabolomics data analysis, researchers are required to fine-tune a multitude of software, settings, and parameters during the bioinformatic processing of raw data^2,3^. It is important to note that these adjustments do not occur in isolation, frequently leading to a cascade of repercussions across various facets of the experiment^4,5^.

In mass spectrometry based metabolomics research, numerous software has been proposed to process the raw data and extract features^6–8^. However, past studies have illuminated significant disparities in the performance of these software tools and algorithmic parameters^9^. For instance, when applied to identical datasets, five distinct software packages yield comparable true positive peaks, yet exhibit variations in the relative quantification of true features. Furthermore, the task of evaluating these software solutions or algorithms is consistently challenging, given the multitude of parameters involved and the inherent uncertainties in non-targeted analysis.

Previous approaches have often relied on the utilization of known compound standards^5^, publicly available datasets^9^ or manually checking^10^ of peak shape to choose software, which might not be reproducible for other researchers. In this context, a “white-box” and reproducible compounds will be helpful to assess software performance or facilitate developmental efforts for metabolomics data analysis.

Proteomics and lipidomics research have used simulation data with a known ground truth^11,12^. LC-MSsim can simulate mzData files with FASTA files as input for proteomics studies^13^.

MSSimulator extended the simulation to MS/MS data in data-dependent acquisition (DDA) mode considering isotope labeling technique^14^. Mspire-Simulator could generate data with post-translational modifications of peptides^15^. JAMSS provides a java based graphical user interface for proteomics data simulation^16^ and Synthedia can cover data-independent acquisition (DIA) mode for proteomics data^12^. The Lipid DDA Simulator can perform in-silico simulation of lipidomics DDA on the Q-Exactive HF based on java^11^. To our knowledge, no R based simulation software has been reported for metabolomics data analysis while R is one of the most popular programming languages for metabolomics data analysis^17^.

Prior efforts in simulating metabolomics data have predominantly concentrated on the compound-level data representation^18,19^. However, real experimental data frequently encompass redundant peaks, offering a glimpse into the intricate nature of metabolomics datasets^20^. Many of these redundant peaks, including isotopologues, adducts, fragmental ions and multiple charged ions, often exhibit correlated intensities with base peaks and are susceptible to the influence of background noise with lower intensity^21^. Such redundant peaks might be useful for downstream annotation analysis^22^. Therefore, in the context of simulation, it becomes imperative to account for the presence of noisy background signals and incorporate the generation of redundant peaks that parallel the characteristics observed in actual experimental data.

Meanwhile, it’s important to note that DDA or DIA modes can result in the loss of MS1 data coverage^23^. A regular workflow in metabolomics always begins with full-scan mode for the relative quantitative analysis, aiming to identify potential biomarkers. Subsequently, MS/MS spectra of selected ions are extracted for annotation purposes. Consequently, the majority of metabolomics software tools are primarily designed to extract MS1 peaks or features^24–27^. In this case, simulation of MS1 data for metabolomics will be helpful to evaluate metabolomics software.

On the other hand, the performance of chromatographic plays a pivotal role, and previous studies have often centered on assessing software based on chromatographic peak quality^28,29^. Nevertheless, achieving consistent peak quality in real samples can be a challenging endeavor considering sample preparation and instrument analysis. Unlike targeted analysis where known ions are optimized, untargeted analysis doesn’t have specific compounds to be optimized. Consequently, issues with peak quality can arise, particularly when dealing with unforeseen compounds. These fluctuations in peak quality might impact the effectiveness of peak picking algorithms. In this context, the inclusion of direct simulation of peak shapes, encompassing characteristics like leading and tailing peaks, can prove highly beneficial. Such simulations would not only provide a more comprehensive evaluation of software performance but also help account for the variability in peak quality encountered in real-world untargeted metabolomics studies.

In this work, we present the ‘mzrtsim’ package in R, designed specifically for the generation of mzML files with the primary aim of evaluating feature extraction or peak picking software. This tool is equipped to produce simulated experimental data that faithfully replicates the issues encountered in real metabolomics datasets. These include the incorporation of redundant peaks and background noise, mirroring the complexity of actual experimental data. Additionally, ‘mzrtsim’ is uniquely capable of simulating chromatographic peak quality issues with reference to known ground truth, providing a comprehensive framework for assessing and enhancing the performance of metabolomics software.

## Methods

The experimental full scan spectrum data from MassBank of North America(MoNA)^30^ for liquid chromatography-mass spectrometry (LC-MS) and the human metabolome database(HMDB)^31^ for gas chromatography-mass spectrometry (GC-MS) was extracted for simulation purpose with redundant peaks. In brief, MoNA database contains 6075 experimental compounds spectra from 1926 unique compounds. 2968 compound spectra from 1114 compounds are collected on high resolution mass spectrometry. Among those spectra, the average peak number is 43.5 and the median peak number is 15. As for HMDB, 10096 experimental spectra are collected from 2295 unique compounds. The average peak number for HMDB is 87.6 and the median peak number is 48. Those experimental data contain not only molecular ions, but also redundant peaks and background noises. Such full scan spectra have been included in the package and could be used for raw data simulation.

For each compound’s spectrum, a response factor (‘rf’) could be set for the base peak to simulate the dynamic range of certain compounds under a predefined baseline level. For example, a compound has two ions with one base peak and another isotopologue of the base peak. The intensity of the isotopologue peak is 5% of the base peaks. Then, the ‘rf’ of 100 with ‘baseline’ 100 for this compound means the base peak of the compound should have 10000 as intensity and 500 for the isotopologue peak. Mass accuracy in ppm can be set for the experimental spectrum with a default value of 5 ppm. To simulate data collected from time of flight mass spectrometry, such value can be set as 20ppm. Another mass accuracy for m/z shift within the sample can also be set to show the fluctuation of m/z within one sample. Such m/z shift is simulated as the m/z shift in ppm multiplying a random number from normal distribution with 0 as mean and 0.5 as standard deviation. Such standard deviation of 0.5 is optimized by comparison of real data from previous studies, which can also be changed by the user as parameters. Meanwhile, a background peak database for LC-MS has been included in the package, which is extracted from solvent blank samples of previous studies^23^. The background peaks can also be simulated by inputting the peak mass to charge ratio.

Each ion in one spectrum will generate its own chromatographic peaks by user defined scan rate, peak height, and peak width. For example, a scan rate of 5 spectra per seconds with 10s as peak width will simulate a peak with 50 spectra. Peak height can be used to simulate the concentration of the same peaks in different samples. For example, peak height of 10 and 15 will multiply the base peak and response factor to show intensity differences of the same peak. Without tailing, each ion in the mass spectra should follow a normal distribution, considering the relative intensity within each spectra. Normal distribution parameters could be set up with mean as ‘baseline’ and standard deviation as ‘baselinesd’ for chromatographic peaks of each ion. The retention time could also be set for each compound. Meanwhile, the tailing factor was introduced to simulate peak quality for certain compounds as it’s widely used in chromatography research^32^. Tailing factor is defined as:

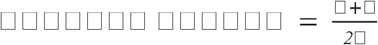

Where a is the peak width at 1/20 peak height for the left part of the peaks and b is the corresponding right part of the peak. Tailing peaks will show a tailing factor larger than 1 and leading peaks will show a tailing factor less than 1 by setting ‘tailingfactor’ in the software.

When the tailing factor is set, two normal distributions with different standard deviations according to the a (¼ peak width) and b (calculated from a and tailing factor) will be generated. Then the whole peak will resemble those two distributions from left to right to show a tailing or leading peak.

With simulation of single spectrum and chromatographic peaks, mzrtsim package^33^ could generate a mzML file through the Spectra package^34^. With control of peak heights for base peaks and retention times, users could generate data sets samples from the same group or different groups. The output files should have mzML files and csv files with mass to charge ratio, retention time, relative intensity and compound name for each peak. Such csv files will be used to evaluate the performance of different software.

mzrtsim package could simulate both GC-MS and LC-MS data with inner databases. The fundamental differences between GC-MS and LC-MS are the ionization process for the compounds and different columns. GC-MS usually needs a hard ionization method to break down the structure of small molecules to generate structure information. However, LC-MS will apply a soft ionization process, which might contain limited information about the structure. Regarding the separation method, GC and LC are all fitting the plate theory. Depending on the separation performance, users can change the scan rate and peak quality for GC-MS or LC-MS raw data simulation. Custom spectra databases can also be used for simulation with MSP file as input.

The major function for mzML files simulation is ‘simmzml’. By setting the compound index in the experimental spectra database, response factor, peaks width and peak height for the base peak of the compounds, retention time of each compound, and file name, mzrtsim package can generate a corresponding mzML file. To change the peak quality, tailing factors of each compound can be changed to generate tailing, leading or normal peaks.

With mzrtsim package (see figure 1), we could benchmark different metabolomics software. We simulate the same data set to compare XCMS^6^, OpenMS^8^, and MZmine^35^. We also test their performance on different types of chromatographic peaks. Then we tested the influence of redundant peaks with lower intensity. The overlapped peaks are defined as peaks with mass to charge ratio shift within 5 ppm and retention time shift within 5 seconds. 1608 matrix ions from previous study^23^ were introduced as background. All the simulated data and codes are available online.

**Figure1.**
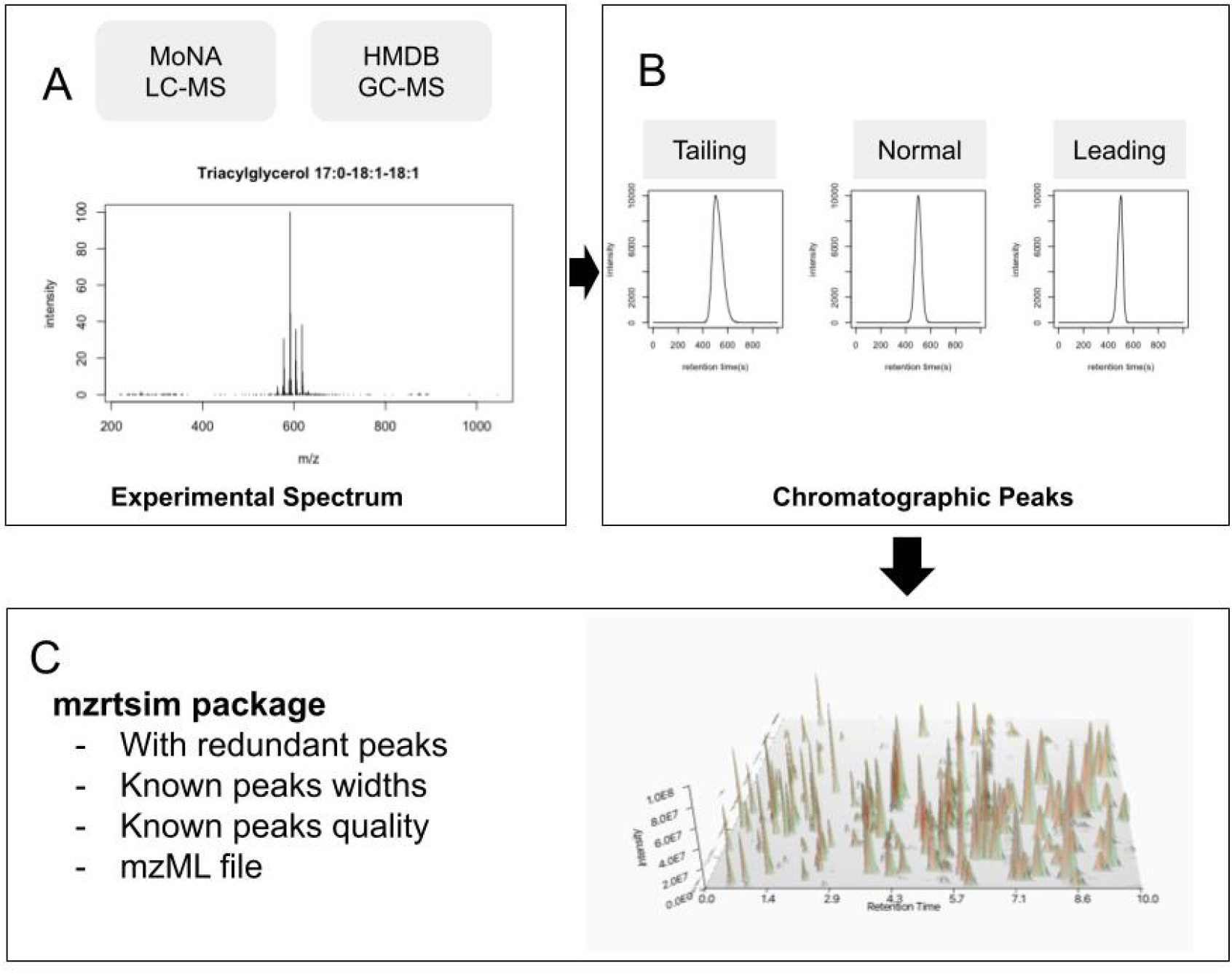
mzML simulation process of mzrtsim package. (A)Experimental spectrum from publicly available databases(MoNA and HMDB) was used for simulation and (B) each ion in the spectrum will generate chromatographic peaks to (C) output the final mzML file with known information for each peak.

## Results and Discussion

### Software comparison for peak picking

We simulated 20 mzML files (10 case samples and 10 control samples) with 100 compounds’ high resolution spectra. Those compounds’ spectra were from MoNA experimental data. The peak widths of base peak were set as 10 seconds and peak heights were set as 5, 10, and 15 for case and control groups at different retention times to make sure around 30 compounds will change. Their retention times were uniformly distributed between 0s and 600s with 3001 MS1 full scan. The response factors for 100 compounds were generated from 100 sampling from 100 and 10000. Those 100 compounds generated 593 detectable chromatographic peaks with a default tailing factor of 1.2. Then peak picking was performed using XCMS (version 4.2.3), OpenMS (pyopenms version 3.0.0), and MZmine (version 4.5) with default settings. To fit the simulated data, the peak width range of XCMS was changed to (5,15) while the other two software’s default settings should cover the simulated peaks. Results showed that those three software found 99 compounds with shared 82 compounds. However, their peak profiles are different. XCMS found 340 peaks and 323 peaks are aligned to real peaks. OpenMS found 564 peaks and 474 peaks are from simulated compounds. MZmine detected 523 peaks, which are all true positive peaks. Noticing only 593 peaks are true peaks, multiple peaks found by the software were aligned to the same true peaks. 4% (XCMS), 16% (OpenMS), and 0% (MZmine) of the detected peaks are false positive peaks and they missed 45% (XCMS), 24% (OpenMS), and 18% (MZmine) true peaks. A closer examination using a Venn diagram (figure 2A) revealed that 272 peaks were shared among the true positives and there were 328(XCMS), 449(OpenMS), and 484(MZmine) true peaks detected. This finding suggests that while the software tools exhibited varying performances in detecting false positive peaks, there was a substantial overlap in their identification of true peaks. Therefore, in practical applications, employing multiple software tools for peak extraction and subsequently focusing on the overlapping peaks is advisable to mitigate the impact of false positives and optimize analysis efficiency. Meanwhile, the false negative peaks’ intensity distribution of those three software are dominated by peaks with lower intensity (figure 2B) for OpenMS and MZmine instead of XCMS. In this case, algorithms for low intensity peaks will improve the coverage of detected peaks for OpenMS and MZmine. For XCMS, 14 out of 18 missing compounds only had one peak and could improve their coverage by considering peaks without redundant peaks.

**Figure 2.**
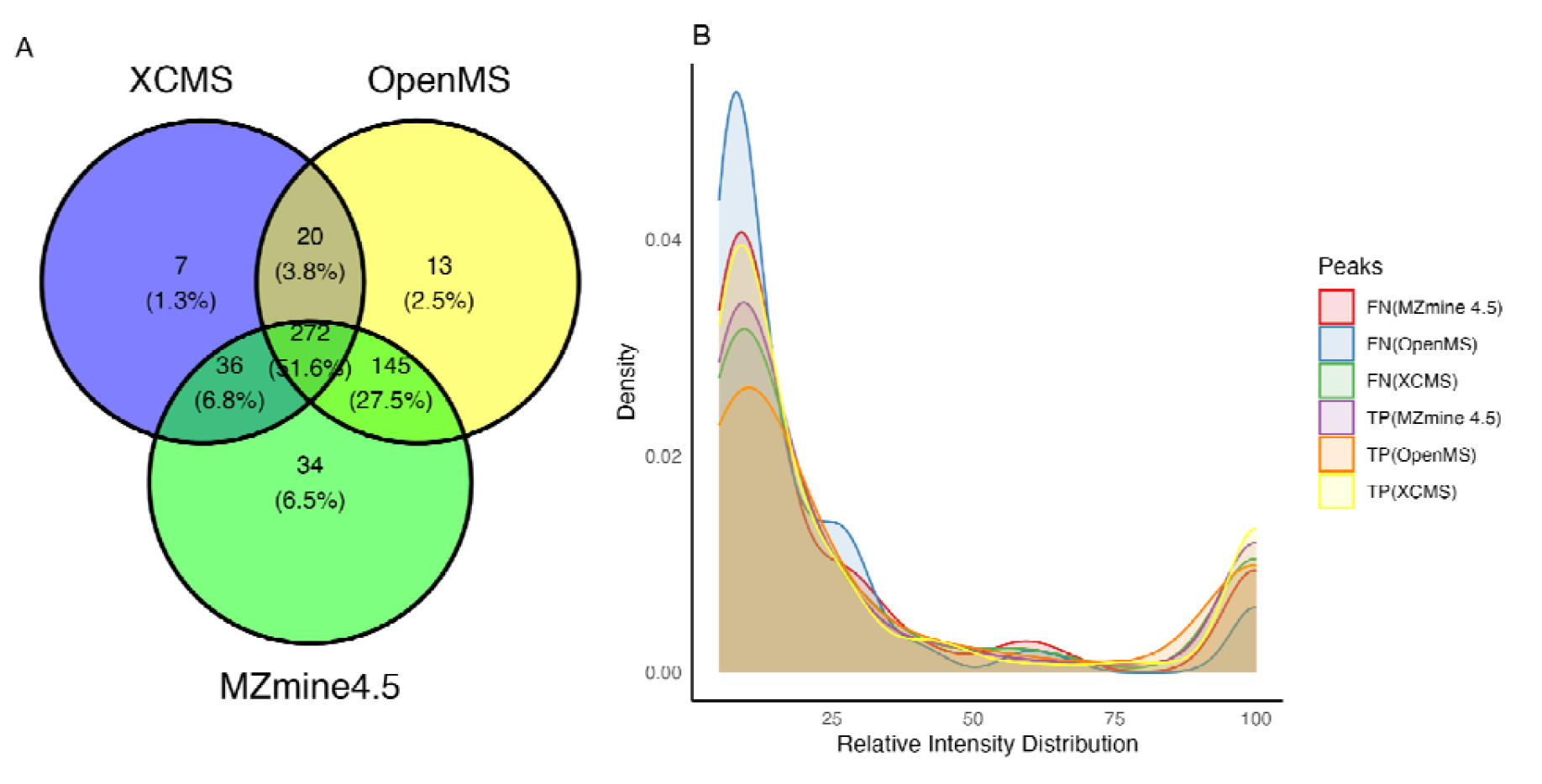
Venn diagram for real detected peaks(left) among XCMS, OpenMS, and MZmine and peak density of relative intensity distribution of simulated peaks (right). TP means true positive peaks, which refer to the real detected peaks. FN means false negative peaks, which refer to the real peaks missing by the software.

### Software comparison for tailing/leading peaks

We also simulate the tailing and leading chromatographic peaks by changing the tailing factor of all peaks. As shown in table 1, different software also displays different sensitivity. XCMS found the minimal numbers of true peaks. OpenMS, on the other hand, finds more false positive peaks in leading peaks than tailing peaks and XCMS shows a worse performance on tailing peaks with more missing peaks and compounds. MZmine also showed a stable performance with least missing peaks. In this case, for data with known leading or tailing peaks, it’s better to use MZmine for peak picking.

**Table 1.**
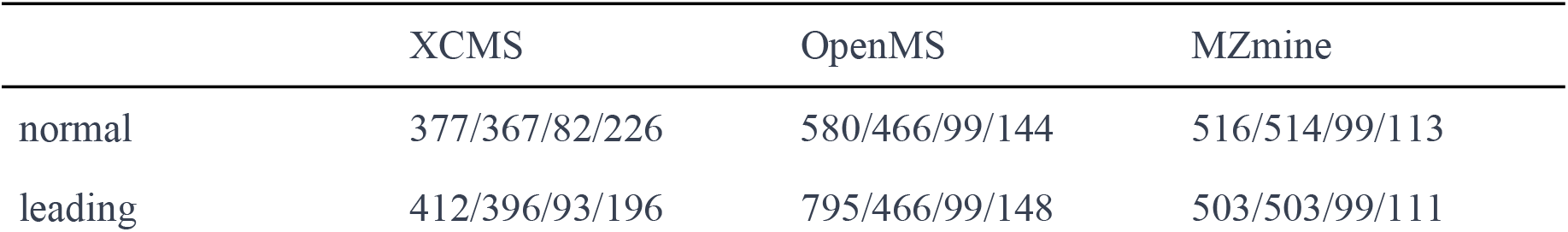
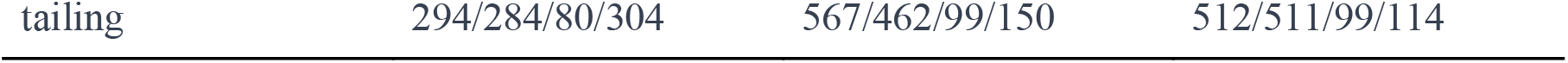
Extracted peaks/True positive peaks/Detected compounds/Missing peaks of XCMS, OpenMS, MZmine peak picking algorithms for 100 simulated compounds with 593 normal (tailing factor 1), leading(tailing factor 0.8) and tailing(tailing factor 1.5) chromatographic peaks.

### Redundant peaks’ influences on statistical analysis

Redundant peaks, characterized by their lack of new biological information and interdependence with other peaks from the same compounds, can affect the efficiency of data analysis. To address this, we conducted experiments involving two groups of samples, each containing known compounds with altered intensities. Importantly, we applied different intensity cutoffs to each set of spectra, with the rationale of reducing the volume of real data for expedited high-throughput analysis.

In a scenario where 30 out of 100 peaks exhibited changes between the two groups of files, we employed intensity cutoffs of 0 and 0.05 to examine their impact on statistical analysis. When the cutoff is set at 0, all peaks derived from the same compounds were retained in the analysis. However, when the cutoff was increased to 0.05, peaks with relative intensities lower than 5% of the base peaks were filtered out. This adjustment resulted in a notable reduction in the typical file size of mzML files, shrinking from 292.3MB to 131.8MB for the same compounds.

Specifically, using a cutoff of 0, we identified 3,746 real peaks from 100 compounds, whereas with a cutoff of 0.05, only 593 peaks remained. Subsequently, employing XCMS for peak extraction, we obtained 2,099 peaks for cutoff 0 and 340 peaks for cutoff 0.05. 1609 real peaks from 28 compounds exhibited changes in files with a cutoff of 0, and 916 peaks were significant according to t-tests with Benjamini-Hochberg procedure^36^ to control false discovery rate, with 19 compounds classified as false positives and 4 compounds classified as false negative. For the cutoff of 0.05, 203 real peaks changed from 28 compounds, and 134 peaks were significant using t-tests with false discovery control, with 2 compounds identified as false positives and 4 compounds classified as false negative. As for OpenMS and MZmine, they found all changed compounds with 1 false positive compound at cutoff 0.05 and also more false positive compounds with cutoff 0 (31 for OpenMS and 14 for MZmine). In this case, the proper cutoff selection will not lose too much information at compound level considering a lower false positive discovery.

### Other potential applications

In addition to comparing various peak-picking algorithms, the simulation of mzML files holds significant potential for enhancing existing algorithms in a broader context within the field of metabolomics. For instance, researchers can utilize this methodology to introduce matrix ions as a form of background noise into their data. Currently, the package includes a matrix ion list from previous published plasma blank samples. This allows for the investigation of how these additional ions may impact subsequent statistical analyses, providing valuable insights into data robustness and the potential need for noise mitigation strategies. Furthermore, when dealing with known compounds with predefined retention times, mzML file simulation offers a unique opportunity to assess the influence of co-elution effects and evaluate the effectiveness of peak alignment algorithms. These insights are crucial for refining data processing pipelines and improving the accuracy of biomarker discovery. Beyond these applications, mzML file simulation can also serve as a valuable tool for validating emerging data analysis methods, such as those based on deep learning techniques and image analysis. By generating controlled datasets with known properties, researchers can rigorously test and fine-tune these innovative approaches, ultimately advancing the state-of-the-art in metabolomics data analysis.

This package also contains limitations. Though we applied experimental data for simulation, the real samples might contain complex matrix effects. The lack of response factor database makes it hard to simulate real samples under certain metabolites profile. Meanwhile, we also left the simulation of retention time to users’ inputs as the elution order of compounds will change on different columns. A compound atlas with corresponding response factor database with levels for certain types of sample will reflect closely to the reality about sample matrix effect. The optimized parameters process of certain peak picking algorithms might also improve the results with simulated ground truth.

## Conclusion

In summary, we have introduced a dedicated raw file simulation tool designed specifically for chromatography coupled with mass spectrometry data. Our initial findings found the presence of false positive peaks and missing true positive peaks. Additionally, our investigation has shed light on the role of redundant peaks, which will not change the information for statistical analysis. The mzrtsim package highlights critical insights into the challenges and opportunities for metabolomics data analysis software development. Such challenges and opportunities included, but are not limited to the optimization for compounds with less peaks, intensity cutoff selection for each spectrum, peak shape adaptation for different column conditions, as well as simulation of various matrix effects.

## Author Contributions

M. Y.: Conceptualization, software development, data curation, visualization, writing-original draft, writing-review and editing; V.P.: writing review and editing, supervision. All authors read, reviewed, and accepted the final manuscript.

## Data Availability Statement

All the data and code associated with the present study can be accessed on Zenodo (https://doi.org/10.5281/zenodo.8299781) and software could be downloaded from GitHub (https://github.com/yufree/mzrtsim)

## Conflicts of Interest

The authors declare no conflict of interests. The authors declare that they have no known competing financial interests or personal relationships that could have appeared to influence the work reported in this paper.

## Notes

https://yufree.github.io/mzrtsim/

### Competing Interest Statement

The authors have declared no competing interest.

### Summary of Updates

Software benchmarks results updates with newer versions of compared software. simulation software updates with new features such as peak height simulation and change snr to response factors.

## References

(1) González-Riano, C.; Dudzik, D.; Garcia, A.; Gil-de-la-Fuente, A.; Gradillas, A.; Godzien, J.; López-Gonzálvez, Á.; Rey-Stolle, F.; Rojo, D.; Ruperez, F. J.; Saiz, J.; Barbas, C. Recent Developments along the Analytical Process for Metabolomics Workflows. Anal. Chem. 2020, 92 (1), 203–226. 10.1021/acs.analchem.9b04553.

(2) Myers, O. D.; Sumner, S. J.; Li, S.; Barnes, S.; Du, X. Detailed Investigation and Comparison of the XCMS and MZmine 2 Chromatogram Construction and Chromatographic Peak Detection Methods for Preprocessing Mass Spectrometry Metabolomics Data. Anal. Chem. 2017, 89 (17), 8689–8695. 10.1021/acs.analchem.7b01069.

(3) Weber, R. J. M.; Lawson, T. N.; Salek, R. M.; Ebbels, T. M. D.; Glen, R. C.; Goodacre, R.; Griffin, J. L.; Haug, K.; Koulman, A.; Moreno, P.; Ralser, M.; Steinbeck, C.; Dunn, W. B.; Viant, M. R. Computational Tools and Workflows in Metabolomics: An International Survey Highlights the Opportunity for Harmonisation through Galaxy. Metabolomics 2017, 13 (2). 10.1007/s11306-016-1147-x.

(4) Liao, J.; Zhang, Y.; Zhang, W.; Zeng, Y.; Zhao, J.; Zhang, J.; Yao, T.; Li, H.; Shen, X.; Wu, G.; Zhang, W. Different Software Processing Affects the Peak Picking and Metabolic Pathway Recognition of Metabolomics Data. J. Chromatogr. A 2023, 1687, 463700. 10.1016/j.chroma.2022.463700.

(5) Li, Z.; Lu, Y.; Guo, Y.; Cao, H.; Wang, Q.; Shui, W. Comprehensive Evaluation of Untargeted Metabolomics Data Processing Software in Feature Detection, Quantification and Discriminating Marker Selection. Anal. Chim. Acta 2018, 1029, 57–. 10.1016/j.aca.2018.05.001.

(6) Domingo-Almenara, X.; Siuzdak, G. Metabolomics Data Processing Using XCMS. In Computational Methods and Data Analysis for Metabolomics; Li, S., Ed.; Methods in Molecular Biology; Springer US: New York, NY, 2020; pp 11–24. 10.1007/978-1-0716-0239-3_2.

(7) Korf, A.; Fouquet, T.; Schmid, R.; Hayen, H.; Hagenhoff, S. Expanding the Kendrick Mass Plot Toolbox in MZmine 2 to Enable Rapid Polymer Characterization in Liquid Chromatography−Mass Spectrometry Data Sets. Anal. Chem. 2020, 92 (1), 628–633. 10.1021/acs.analchem.9b03863.

(8) Röst, H. L.; Sachsenberg, T.; Aiche, S.; Bielow, C.; Weisser, H.; Aicheler, F.; Andreotti, S.; Ehrlich, H.-C.; Gutenbrunner, P.; Kenar, E.; Liang, X.; Nahnsen, S.; Nilse, L.; Pfeuffer, J.; Rosenberger, G.; Rurik, M.; Schmitt, U.; Veit, J.; Walzer, M.; Wojnar, D.; Wolski, W. E.; Schilling, O.; Choudhary, J. S.; Malmström, L.; Aebersold, R.; Reinert, K.; Kohlbacher, O. OpenMS: A Flexible Open-Source Software Platform for Mass Spectrometry Data Analysis. Nat. Methods 2016, 13 (9), 741–748. 10.1038/nmeth.3959.

(9) El Abiead, Y.; Milford, M.; Schoeny, H.; Rusz, M.; Salek, R. M.; Koellensperger, G. Power of mzRAPP-Based Performance Assessments in MS1-Based Nontargeted Feature Detection. Anal. Chem. 2022, 94 (24), 8588–8595. 10.1021/acs.analchem.1c05270.

(10) Chetnik, K.; Petrick, L.; Pandey, G. MetaClean: A Machine Learning-Based Classifier for Reduced False Positive Peak Detection in Untargeted LC–MS Metabolomics Data. Metabolomics Off. J. Metabolomic Soc. 2020, 16 (11), 117. 10.1007/s11306-020-01738-3.

(11) Hutchins, P. D.; Russell, J. D.; Coon, J. J. Accelerating Lipidomic Method Development through in Silico Simulation. Anal. Chem. 2019, 91 (15), 9698–9706. 10.1021/acs.analchem.9b01234.

(12) Leeming, M. G.; Ang, C.-S.; Nie, S.; Varshney, S.; Williamson, N. A. Simulation of Mass Spectrometry-Based Proteomics Data with Synthedia. Bioinforma. Adv. 2023, 3 (1), vbac096. 10.1093/bioadv/vbac096.

(13) Schulz-Trieglaff, O.; Pfeifer, N.; Gröpl, C.; Kohlbacher, O.; Reinert, K. LC-MSsim--a Simulation Software for Liquid Chromatography Mass Spectrometry Data. BMC Bioinformatics 2008, 9, 423. 10.1186/1471-2105-9-423.

(14) Bielow, C.; Aiche, S.; Andreotti, S.; Reinert, K. MSSimulator: Simulation of Mass Spectrometry Data. J. Proteome Res. 2011, 10 (7), 2922–2929. 10.1021/pr200155f.

(15) Noyce, A. B.; Smith, R.; Dalgleish, J.; Taylor, R. M.; Erb, K. C.; Okuda, N.; Prince, J. T. Mspire-Simulator: LC-MS Shotgun Proteomic Simulator for Creating Realistic Gold Standard Data. J. Proteome Res. 2013, 12 (12), 5742–5749. 10.1021/pr400727e.

(16) Smith, R.; Prince, J. T. JAMSS: Proteomics Mass Spectrometry Simulation in Java. Bioinformatics 2015, 31 (5), 791–793. 10.1093/bioinformatics/btu729.

(17) Shen, X.; Yan, H.; Wang, C.; Gao, P.; Johnson, C. H.; Snyder, M. P. TidyMass an Object-Oriented Reproducible Analysis Framework for LC–MS Data. Nat. Commun. 2022, 13 (1), 4365. 10.1038/s41467-022-32155-w.

(18) Manimaran, S.; Selby, H. M.; Okrah, K.; Ruberman, C.; Leek, J. T.; Quackenbush, J.; Haibe-Kains, B.; Bravo, H. C.; Johnson, W. E. BatchQC: Interactive Software for Evaluating Sample and Batch Effects in Genomic Data. Bioinformatics 2016, 32 (24), 3836–3838. 10.1093/bioinformatics/btw538.

(19) Mendes, P.; Camacho, D.; de la Fuente, A. Modelling and Simulation for Metabolomics Data Analysis. Biochem. Soc. Trans. 2005, 33 (6), 1427–1429. 10.1042/BST0331427.

(20) Mahieu, N. G.; Patti, G. J. Systems-Level Annotation of a Metabolomics Data Set Reduces 25 □ 000 Features to Fewer than 1000 Unique Metabolites. Anal. Chem. 2017, 89 (19), 10397–10406. 10.1021/acs.analchem.7b02380.

(21) Yu, M.; Olkowicz, M.; Pawliszyn, J. Structure/Reaction Directed Analysis for LC-MS Based Untargeted Analysis. Anal. Chim. Acta 2019, 1050, 24–. 10.1016/j.aca.2018.10.062.

(22) Chen, L.; Lu, W.; Wang, L.; Xing, X.; Chen, Z.; Teng, X.; Zeng, X.; Muscarella, A. D.; Shen, Y.; Cowan, A.; McReynolds, M. R.; Kennedy, B. J.; Lato, A. M.; Campagna, S. R.; Singh, M.; Rabinowitz, J. D. Metabolite Discovery through Global Annotation of Untargeted Metabolomics Data. Nat. Methods 2021, 18 (11), 1377–1385. 10.1038/s41592-021-01303-3.

(23) Yu, M.; Dolios, G.; Petrick, L. Reproducible Untargeted Metabolomics Workflow for Exhaustive MS2 Data Acquisition of MS1 Features. J. Cheminformatics 2022, 14 (1), 6. 10.1186/s13321-022-00586-8.

(24) Libiseller, G.; Dvorzak, M.; Kleb, U.; Gander, E.; Eisenberg, T.; Madeo, F.; Neumann, S.; Trausinger, G.; Sinner, F.; Pieber, T.; Magnes, C. IPO: A Tool for Automated Optimization of XCMS Parameters. BMC Bioinformatics 2015, 16, 118. 10.1186/s12859-015-0562-8.

(25) Delabriere, A.; Warmer, P.; Brennsteiner, V.; Zamboni, N. SLAW: A Scalable and Self-Optimizing Processing Workflow for Untargeted LC-MS. Anal. Chem. 2021, 93 (45), 15024–15032. 10.1021/acs.analchem.1c02687.

(26) McLean, C.; Kujawinski, E. B. AutoTuner: High Fidelity and Robust Parameter Selection for Metabolomics Data Processing. Anal. Chem. 2020, 92 (8), 5724–5732. 10.1021/acs.analchem.9b04804.

(27) Ni, Y.; Su, M.; Qiu, Y.; Jia, W.; Du, X. ADAP-GC 3.0: Improved Peak Detection and Deconvolution of Co-Eluting Metabolites from GC/TOF-MS Data for Metabolomics Studies. Anal. Chem. 2016, 88 (17), 8802–8811. 10.1021/acs.analchem.6b02222.

(28) Gloaguen, Y.; Kirwan, J. A.; Beule, D. Deep Learning-Assisted Peak Curation for Large-Scale LC-MS Metabolomics. Anal. Chem. 2022, 94 (12), 4930–4937. 10.1021/acs.analchem.1c02220.

(29) Guo, J.; Shen, S.; Huan, T. Paramounter: Direct Measurement of Universal Parameters To Process Metabolomics Data in a “White Box.” Anal. Chem. 2022. 10.1021/acs.analchem.1c04758.

(30) Horai, H.; Arita, M.; Kanaya, S.; Nihei, Y.; Ikeda, T.; Suwa, K.; Ojima, Y.; Tanaka, K.; Tanaka, S.; Aoshima, K.; Oda, Y.; Kakazu, Y.; Kusano, M.; Tohge, T.; Matsuda, F.; Sawada, Y.; Hirai, M. Y.; Nakanishi, H.; Ikeda, K.; Akimoto, N.; Maoka, T.; Takahashi, H.; Ara, T.; Sakurai, N.; Suzuki, H.; Shibata, D.; Neumann, S.; Iida, T.; Tanaka, K.; Funatsu, K.; Matsuura, F.; Soga, T.; Taguchi, R.; Saito, K.; Nishioka, T. MassBank: A Public Repository for Sharing Mass Spectral Data for Life Sciences. J. Mass Spectrom. 2010, 45 (7), 703–714. 10.1002/jms.1777.

(31) Wishart, D. S.; Feunang, Y. D.; Marcu, A.; Guo, A. C.; Liang, K.; Vázquez-Fresno, R.; Sajed, T.; Johnson, D.; Li, C.; Karu, N.; Sayeeda, Z.; Lo, E.; Assempour, N.; Berjanskii, M.; Singhal, S.; Arndt, D.; Liang, Y.; Badran, H.; Grant, J.; Serra-Cayuela, A.; Liu, Y.; Mandal, R.; Neveu, V.; Pon, A.; Knox, C.; Wilson, M.; Manach, C.; Scalbert, A. HMDB 4.0: The Human Metabolome Database for 2018. Nucleic Acids Res. 2018, 46 (D1), D608–D617. 10.1093/nar/gkx1089.

(32) Petersson, P.; Forssen, P.; Edström, L.; Samie, F.; Tatterton, S.; Clarke, A.; Fornstedt, T. Why Ultra High Performance Liquid Chromatography Produces More Tailing Peaks than High Performance Liquid Chromatography, Why It Does Not Matter and How It Can Be Addressed. J. Chromatogr. A 2011, 1218 (39), 6914–6921. 10.1016/j.chroma.2011.08.018.

(33) Yu, M.; Roszkowska, A.; Pawliszyn, J. Simulation-Based Comprehensive Study of Batch Effects in Metabolomics Studies. bioRxiv December 17, 2019, p 2019.12.16.878637. 10.1101/2019.12.16.878637.

(34) Rainer, J.; Vicini, A.; Salzer, L.; Stanstrup, J.; Badia, J. M.; Neumann, S.; Stravs, M. A.; Verri Hernandes, V.; Gatto, L.; Gibb, S.; Witting, M. A Modular and Expandable Ecosystem for Metabolomics Data Annotation in R. Metabolites 2022, 12 (2), 173. 10.3390/metabo12020173.

(35) Schmid, R.; Heuckeroth, S.; Korf, A.; Smirnov, A.; Myers, O.; Dyrlund, T. S.; Bushuiev, R.; Murray, K. J.; Hoffmann, N.; Lu, M.; Sarvepalli, A.; Zhang, Z.; Fleischauer, M.; Dührkop, K.; Wesner, M.; Hoogstra, S. J.; Rudt, E.; Mokshyna, O.; Brungs, C.; Ponomarov, K.; Mutabdžija, L.; Damiani, T.; Pudney, C. J.; Earll, M.; Helmer, P. O.; Fallon, T. R.; Schulze, T.; Rivas-Ubach, A.; Bilbao, A.; Richter, H.; Nothias, L.-F.; Wang, M.; Orešič, M.; Weng, J.-K.; Böcker, S.; Jeibmann, A.; Hayen, H.; Karst, U.; Dorrestein, P. C.; Petras, D.; Du, X.; Pluskal, T. Integrative Analysis of Multimodal Mass Spectrometry Data in MZmine 3. Nat. Biotechnol. 2023, 41 (4), 447–449. 10.1038/s41587-023-01690-2.

(36) Benjamini, Y.; Hochberg, Y. Controlling the False Discovery Rate: A Practical and Powerful Approach to Multiple Testing. J. R. Stat. Soc. Ser. B Methodol. 1995, 57 (1), 289–300.

